# Long read sequencing provides an insight into plasmids found among carbapenemase producing *Enterobacterales* from hospitals in the United Kingdom during 2021 to 2023

**DOI:** 10.1101/2024.03.19.585710

**Authors:** Jane F Turton, Jack A Turton

**Author notes:** Sequences were deposited on GenBank under project <u>PRJNA1010831</u>.

## Abstract

107 isolates of *Enterobacterales* consisting of *Klebsiella pneumoniae* (n=90), *Escherichia coli* (n=7), *Enterobacter cloacae* complex (n=6), *Klebsiella oxytoca* complex (n=3) and *Citrobacter freundi* (n=1) and additionally an isolate of *Acinetobacter baumannii* carrying genes encoding NDM (NDM-1, NDM-5 and NDM-14), KPC (KPC-2 and KPC-3), OXA-48-like (OXA-48, OXA-181 and OXA-232) and IMP (IMP-1 and IMP-4) carbapenemases were sequenced using q20 nanopore chemistry to provide complete/near-complete assemblies and relevant plasmids compared. Investigation of potential ‘plasmid outbreaks’ in individual hospitals among isolates of different types and species revealed a mixed situation with some isolates carrying similar plasmids, but with segments missing/added and some plasmids that were clearly distinct.

While most plasmids carrying *bla*_OXA-48_ were typical IncL plasmids of approximately 60 kb that are widely described, there was some variation among these. One isolate carried an IncR plasmid that had only limited homology with the others. Identical 51,479 bp ColKP3/IncX3 plasmids carrying *bla*_OXA-181_ were found from isolates from different hospitals that exactly matched those on GenBank from other countries. In other isolates *bla*_OXA-181_ was carried on IncFII plasmids. *bla*_OXA-232_ was found in highly conserved small ColKP3 plasmids that matched those found in other countries and continents. These observations highlight the importance of understanding the wider distribution of plasmids of concern.

IncHI2/IncHI2A plasmids were important vehicles for carbapenemase genes and were found with *bla*_KPC-2_, *bla*_IMP-1_, *bla*_IMP-4_ or *bla*_NDM-1_, sometimes with the colistin resistance gene *mcr-9* in addition. Representatives of *K. pneumoniae* sequence type (ST) 147 from seven hospitals carried IncFIB(pNDM-Mar)/IncHI1B(pNDM-MAR) hybrid virulence resistance plasmids of 325 to 352 kb that combined *bla*_NDM-5_ and other resistance genes with genes found in virulence plasmids. A similar plasmid was also found in an isolate of *K. pneumoniae* ST1558 and has been described in representatives of ST383.

Nanopore sequencing has been instrumental in improving our knowledge of plasmids carrying carbapenemase genes leading to a better understanding of their epidemiology.

## Introduction

Carbapenems have been considered ‘last-resort’ antibiotics for the treatment of infections caused by members of the *Enterobacterales* belonging to genera such as *Klebsiella, Enterobacter, Escherichia* and *Citrobacter*. However, the dramatic rise in incidence of carbapenemase enzymes, especially belonging to the NDM, OXA-48-like, KPC, IMP and VIM families, that can degrade these antibiotics among these organisms seriously threatens their effectiveness [1]. The genes encoding these carbapenemases are mostly carried in plasmids, but in many instances, these remain uncharacterised.

The advent of q20 high accuracy nanopore sequencing [2] has facilitated complete assembly of bacterial chromosome and their plasmids, providing an opportunity to investigate and describe plasmids, including those carrying carbapenemase genes. This not only provides a better understanding of their genetic environment but enables an assessment of the extent to which plasmids or other elements carrying carbapenemase genes may be shared between organisms of different species from the same and different hospitals. Here we describe sequences from isolates of *Klebsiella pneumoniae, Escherichia coli, Enterobacter cloacae* complex, *Klebsiella oxytoca* complex, *Citrobacter freundi* and additionally an isolate of *Acinetobacter baumannii* carrying a variety of carbapenemase genes encoding NDM (NDM-1, NDM-5 and NDM-14), KPC (KPC-2 and KPC-3), OXA-48-like (OXA-48, OXA-181 and OXA-232) and IMP (IMP-1 and IMP-4) carbapenemases. In order to answer various questions around potentially shared plasmids and around suitability of the method for typing of *Klebsiella pneumoniae*, there is a focus on isolates from particular hospitals and on particular types, especially *K. pneumoniae* Sequence Types (ST) 147, 307 and 20, but the 108 isolates included consisted of many types and were from a total of 30 hospital laboratories. This work provides an insight into plasmids circulating among *Enterobacterales* in UK hospitals between 2021 and 2023.

## Methods

Isolates were referred to the UK reference laboratory for typing from hospitals in the UK and were typed by Variable Number Tandem Repeat (VNTR) analysis at 11 loci (*Klebsiella pneumoniae sensu stricto*) or pulsed-field gel electrophoresis of XbaI-digested genomic DNA (all other species). They were labelled by an organism code (KP for *K. pneumoniae*, EB for *E. cloacae* complex, ES for *E. coli*, CF for *C. freundii*, KO for *K. oxytoca* complex and AB for *Acinetobacter baumannii*) followed by a unique number for that organism, hospital code, the week and year of receipt, their sequence type and the carbapenemase genes that they carried (e.g., KP2_L3_37.21_ST147_NDM5). Hospitals were labelled according to region and number within that region (L, London; SE, South East England; SW, South West England; WM, West Midlands; EM, East Midlands; NE, North East London; NW, North West England; YH, Yorkshire and Humber). Patient duplicates (i.e. more than one isolate of the same type from the same patient) were excluded. All isolates are listed in Table S1.

### Nanopore sequencing

DNA from growth from single colony picks was extracted using a GeneJet genomic DNA kit (ThermoFisher, UK) and sequenced on R10.4.1 flow cells on a minION Mk1C (or gridION for run q20run8) following library preparation using the rapid barcoding kit SQK-RBK114.24 (Oxford Nanopore Technologies, UK). Library preparation was carried out following the protocol provided by Oxford Nanopore Technologies, with concentration of the pooled barcoded libraries using ‘clumping buffer’ as described previously [3] (rrp8, rrp9, q20runs 3-7) or Ampure beads (q20runs 8-16). Basecalling was by the high accuracy basecalling option with barcode trimming. MinKNOW versions were 22.10.7 for q20 run 2 (rrp9), 22.12.5 (bionic) for q20 runs 5, 7, 8, 10, 13 and 14 and 23.04.5 (bionic) for q20 runs 15 and 16. A speed of 260 bp/s was used unless only a 400 bp/s option was available. Assemblies were done using flye 2.9.1-b1780 [4] and medaka 1.7.2 (https://github.com/nanoporetech/medaka). Resistance genes, plasmid replicon types, plasmid sequence types (pST) and MLST types were sought using ResFinder 4.3.3, PlasmidFinder 2.1, pMLST 2.0 and MLST 2.0 on the Center for Genomic Epidemiology website (https://cge.food.dtu.dk/) [5, 6, 7].

### Resistance genes

Resistance genes were as detected on ResFinder. Where there was no exact match, the closest allele(s) are provided. As a guide, the antibiotic classes to which the most common acquired genes found in this study confer resistance are summarised in Table 1, but fuller information is available on ResFinder (http://genepi.food.dtu.dk/resfinder).

**Table 1.** Guide to antibiotic resistance classes to which acquired resistance genes found in this study confer resistance. Fuller information is available from ResFinder (http://genepi.food.dtu.dk/resfinder).

| Resistance class | Resistance genes beginning |
| --- | --- |
| aminoglycoside | *aac*, *aad*, *ant*, *aph*, *armA*, *rmt* |
| quinolone | *qnr* |
| Beta-lactam | *bla*<sub>KPC</sub>, *bla*<sub>CTX-M</sub>, *bla*<sub>TEM</sub>, *bla*<sub>SHV</sub>, *bla*<sub>IMP</sub>, *bla*<sub>NDM</sub>, *bla*<sub>ACT</sub>, *bla*<sub>CMY</sub>, *bla*<sub>OXA</sub>, *bla*<sub>ADC</sub> |
| folate pathway antagonist | *sul*, *dfrA* |
| tetracycline | *tet* |
| Streptogramins | *msr* |
| macrolides | *ere*, *erm*, *mph* |
| polymyxin | *mcr* |
| amphenicol | *cat* |

### Sequences

Sequences were deposited on GenBank under project <u>PRJNA1010831</u>.

## Results and Discussion

### Isolates from hospitals querying ‘plasmid outbreaks’

Many of these isolates were referred by hospitals concerned that they may be experiencing plasmid outbreaks and often consisted of sets containing multiple species of *Enterobacterales*. Here we describe the results of investigation of three such sets.

#### a) Isolates from hospital WM3 carrying *bla*_KPC-2_

Thirteen isolates, all carrying *bla*_KPC-2_, were sequenced, and carried a mixture of plasmid types. Four isolates (EB4_WM3_44.22_NV1_KPC2, KO1_WM3_47.22_ST183_KPC2, KO3_WM3_07.23_ST77_KPC2 and KP111_WM3_19.23_ST846_KPC2) belonging to three different species (*Enterobacter hormaechei, Klebsiella michiganensis* and *K. pneumoniae*) and all of different types carried an IncN plasmid of pST15 of 63-69 kb carrying *bla*_KPC-2_, *aph(6)-Id, aph(3”)-Ib, bla*_TEM-1B_, *qnrB2, sul1* and *dfrA19* (e.g. <u>CP141849</u>) (Fig. 1). On GenBank the most similar plasmid was also in an isolate (S11_16) of *E. hormaechei* from the UK (<u>CP035387</u>) collected in 2016. However, other isolates (EB6_WM3_49.22_ST66_KPC2 and ES6_WM3_50.22_ST95_KPC2) carried an IncHI2/IncHI2A plasmid of approximately 300 kb with *bla*_KPC-2_, *aadA2b, ant(2”)-Ia, bla*_CTX-M-9_, *qnrA1, sul1, tet(A), dfrA16* (e.g. <u>CP133857</u>); a further isolate (KP71_WM3_51.22_ST462_KPC2) carried a larger IncHI2/IncHI2A plasmid (461 kb) containing those same elements. Two isolates appeared to carry these elements chromosomally (KP66_WM3_49.22_ST307_KPC2 and ES7_WM3_51.22_ST5295_KPC2); this was the case for two separate runs carried out on these isolates. At least two other plasmid types were also detected among the set, a 220 kb IncHI1B(pNDM-MAR)/ IncFIB(pNDM-Mar) plasmid carrying *bla*_KPC-2_, *aph(3”)-Ib, aph(6)-Id, bla*_TEM-1B_, *qnrB2, sul1* and *dfrA19*, (in KP64_WM3_49.22_ST307_KPC2) (CP141558) and a 270 kb IncFIB(pQil) plasmid (in KP49_WM3_45.22_ST258_KPC2).

**Fig. 1.**
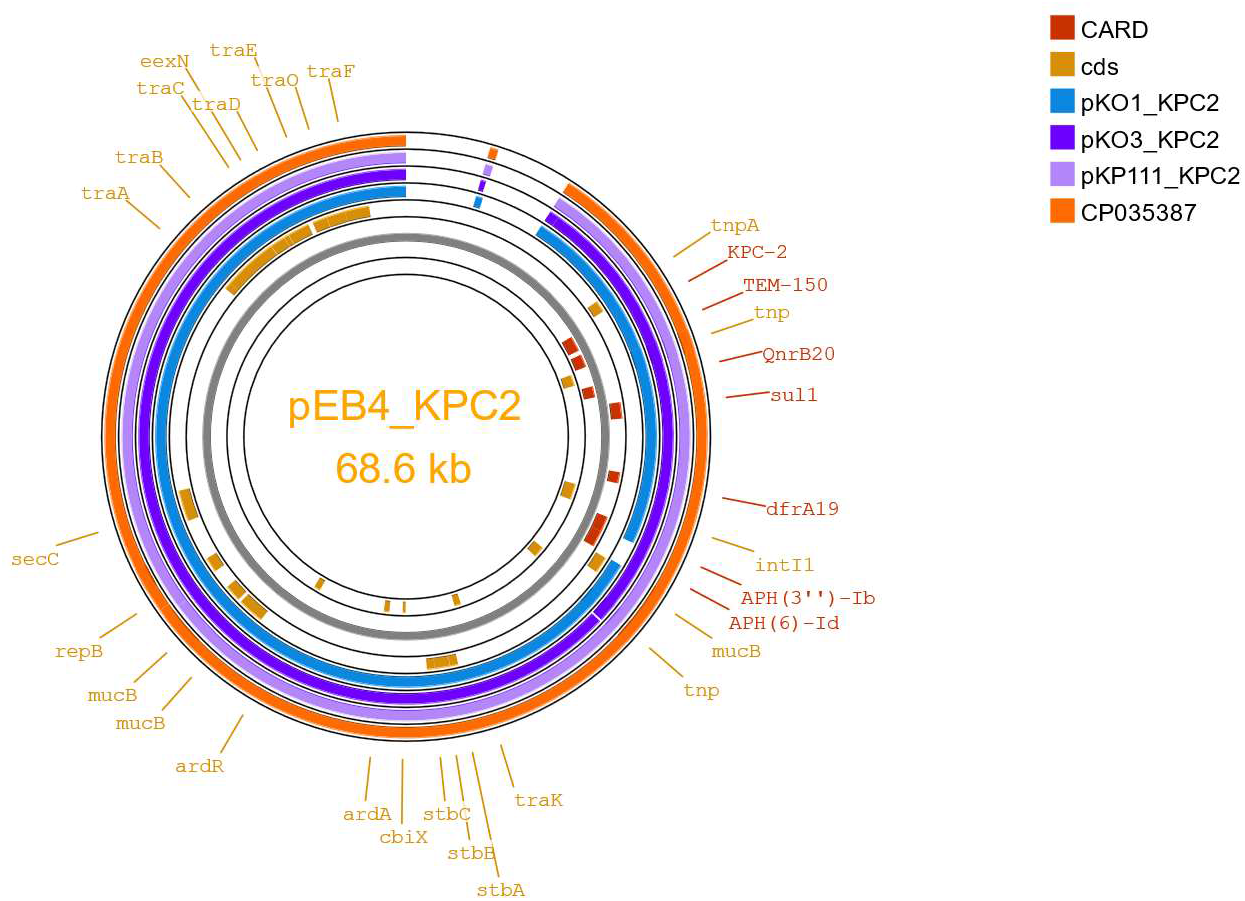
BLAST comparison of IncN plasmids of pST15 carrying *bla*_KPC-2_ from isolates from hospital WM3 and with that of *E. hormaechei* S11_16 (<u>CP035387)</u>. Plasmids were compared with that of isolate EB4_WM3_44.22_NV1_KPC2 (pEB4_KPC2). Regions of homology are shown in colour. Plasmids were from isolates of three different species (EB, *Enterobacter cloacae* complex; KO, *Klebsiella oxytoca* group and KP, *Klebsiella pneumoniae*) and were compared by Proksee [8]. Resistance genes were identified by the Comprehensive Antibiotic Resistance Database (CARD) on Proksee. NV1 refers to a novel sequence type of the *Enterobacter cloacae* complex of allelic profile 59,9,12,9,242,37,6. Only named coding sequences (cds) are shown.

#### b) Isolates from hospital L5 carrying *bla*_NDM_

Many of the isolates from this investigation were clonally related isolates of *K. pneumoniae* ST20 carrying *bla*_NDM-1_. These carried an IncHI2/ IncHI2A plasmid of approximately 340 kb carrying *bla*_NDM-1_, *aadA5, aac(6’)-Ib-cr, aac(6’)-IIc, bla*_DHA-1_, *bla*_OXA-1_, *ere(A), mph(A), catB3, qnrB4, sul1, dfrA17* (e.g. <u>CP141854</u> from KP34_L5_41.22_ST20_NDM1). However, there were also isolates of *K. pneumoniae* of ST16 carrying IncX3 plasmids of 46 kb carrying *bla* _NDM-5_ with no other resistance elements detected; these isolates also carried *bla*_OXA-232_ in a relatively small (12.3 kb) ColKP3 plasmid (KP21_L5_33.22_ST16_NDM5_OXA232 and KP24_L5_34.22_ST16_NDM5_OXA232). There were also two further types of *K. pneumoniae* (ST147 and ST3096) carrying *bla*_NDM-5_, each with further distinct plasmids, the two representatives of ST147 carrying a 156.5 kb IncFII/IncR plasmid with *bla*_NDM-5_, *aadA2, aadA1, armA, rmtB, bla*_CTX-M-15_, *bla*_TEM-1B_, *msr(E), ere(A), mph(A), mph(E), erm(B), cmlA1, qnrB1, ARR-2, sul1* and *dfrA12* (KP104_L5_18.23_ST147_NDM5_OXA232) and a 343.0 kb IncFIB(pNDM-Mar)/IncHI1B(pNDM-MAR) hybrid virulence/resistance plasmid (carrying *bla*_NDM-5_, *aadA1, aph(3’)-VI, aac(6’)-Ib, bla*_CTX-M-15_, *bla*_TEM-1C/1B_, *bla*_OXA-9_, *mph(A)*, truncated *catA1, qnrS1, sul1, sul2, dfrA5* together with various genes associated with virulence plasmids (*rmpA*/*A2, terABCDEWXYZ, iucABCD,iutA*, hemin, *shiF*, lysozyme inhibitor, *cobW*)) (KP124_L5_21.23_ST147_NDM5) (<u>CP137392</u>), respectively, while the isolate of ST3096 (KP52_L5_46.22_ST3096_NDM5_OXA48) carried a 143 kb IncC plasmid with *bla*_NDM-5_, *aac(6’)-Ib3, bla*_CMY-23_ and *sul1* (<u>CP141561</u>); this isolate also carried a 78.9 kb IncL plasmid with *bla*_OXA-48_, *aac(6’)-Ib-cr, bla*_TEM-1D_, *bla*_OXA-1_, truncated *catB3* and *qnrS1* (<u>CP141563</u>).

#### c) Isolates from hospital WM2 carrying *bla*_OXA-48_ and/or *bla*_NDM-_5

A set of seven isolates belonging to four different genera were sequenced, five of which carried *bla*_OXA-48_. Four of the five carried IncL plasmids (CF2_WM2_48.22_ST162_OXA48, EB5_WM2_48.22_ST108_OXA48, ES5_WM2_48.22_ST1607_OXA48 and KP58_WM2_48.22_ST147_NDM5_OXA48) (<u>CP133849</u>, <u>CP133863</u>, <u>CP133873</u> and <u>CP137377</u>, respectively) only one of which carried other resistance determinants in addition to *bla*_OXA-48_ (*aph(3”)-Ib, aph(6)-Id, bla*_CTX-M-14b_ in <u>CP137377</u>). The remaining isolate (KP59_WM2_48.22_ST395_OXA48) carried a 54.8 kb IncR plasmid that carried *catA1* in addition to *bla*_OXA-48_. Three isolates carried *bla*_NDM-5_, one in a 459.9 kb IncFIB(pNDM-Mar)/IncHI1B(pNDM-MAR) hybrid virulence/resistance plasmid (carrying *bla*_NDM-5_, *aac(6’)-Ib-cr, aadA1, aph(3’)-VI, aac(6’)-Ib, bla*_CTX-M-15_, *bla*_TEM-1C/1B_, *bla*_OXA-9_, *mph(A)*, truncated *catA1, qnrS1, sul1* and *dfrA5* together with virulence plasmid elements *rmpA*/*A2, terABCDEWXYZ, iucABCD, iutA*, hemin, *shiF*, lysozyme inhibitor and *cobW*) (<u>CP137375</u>). The remaining isolates carrying *bla*_NDM-5_ were both representatives of *E. coli* ST167 with multi-replicon IncF (IncFIA /IncFIB(AP001918)/IncFII ) *bla*_NDM-5_ plasmids (e.g. <u>CP133853</u>) widely found in this organism [9, 10, 11]; they were highly similar to one another, one of 143.5 kb and the other of 141.9 kb, both with *bla*_NDM-5_, *aac(6’)-Ib-cr, aadA2, sul1, dfrA12, sitABCD, tet(A), bla*_CTX-M-15_, *bla*_OXA-1_, *mph(A)*, truncated *qacE* and *catB3* (ES3_WM2_48.22_ST167_NDM5 and ES4_WM2_48.22_ST167_NDM5).

In all three cases, a mixed scenario was observed with some isolates carrying highly similar plasmids, while there were also clear examples where the plasmids were distinct.

### OXA-48 plasmids

21 isolates in the set from eight hospital groups carried *bla*_OXA-48_, the vast majority in IncL plasmids of approximately 64 kb that contained no other resistance genes (e.g. <u>CP133849</u>, <u>CP133873</u>, <u>CP133846</u>). These have a large number of matches on GenBank with these plasmids being widely distributed. In some cases the IncL plasmid was larger (e.g. <u>CP133863</u> (130 kb), <u>CP137356</u> (72.2 kb)) (Fig. 2). The latter included an extra segment that carried *aph(6)-Id, aph(3”)-Ib, aph(3’)-Vib* and *bla*_CTX-M-14b_. There were 5 examples of isolates with a 61.5 kb IncM1 *bla*_OXA-48_ plasmid with no other resistance genes; these were all representatives of *K. pneumoniae* ST307 from the same hospital related in space and time (KP44 to KP48 inclusive all followed by _NW4_44.22_ST307_OXA48) (e.g. <u>CP141564</u> in KP44_NW4_44.22_ST307_OXA48). KP59_WM2_48.22_ST395_OXA48 carried a 54.8 kb IncR *bla*_OXA-48_ plasmid also containing *catA1*, while KP17_NW3_25.22_ST395_OXA48 carried a 78.1 kb IncM2 *bla*_OXA-48_ plasmid also carrying *msr(E), armA, mph(E), bla*_TEM-1B_ resistance genes (<u>CP133747</u>). The IncR plasmid (pKP59_OXA48) shown in orange in Fig. 2) only had limited homology with the IncL, IncM1 and IncM2 plasmids (Fig. 2).

**Fig. 2.**
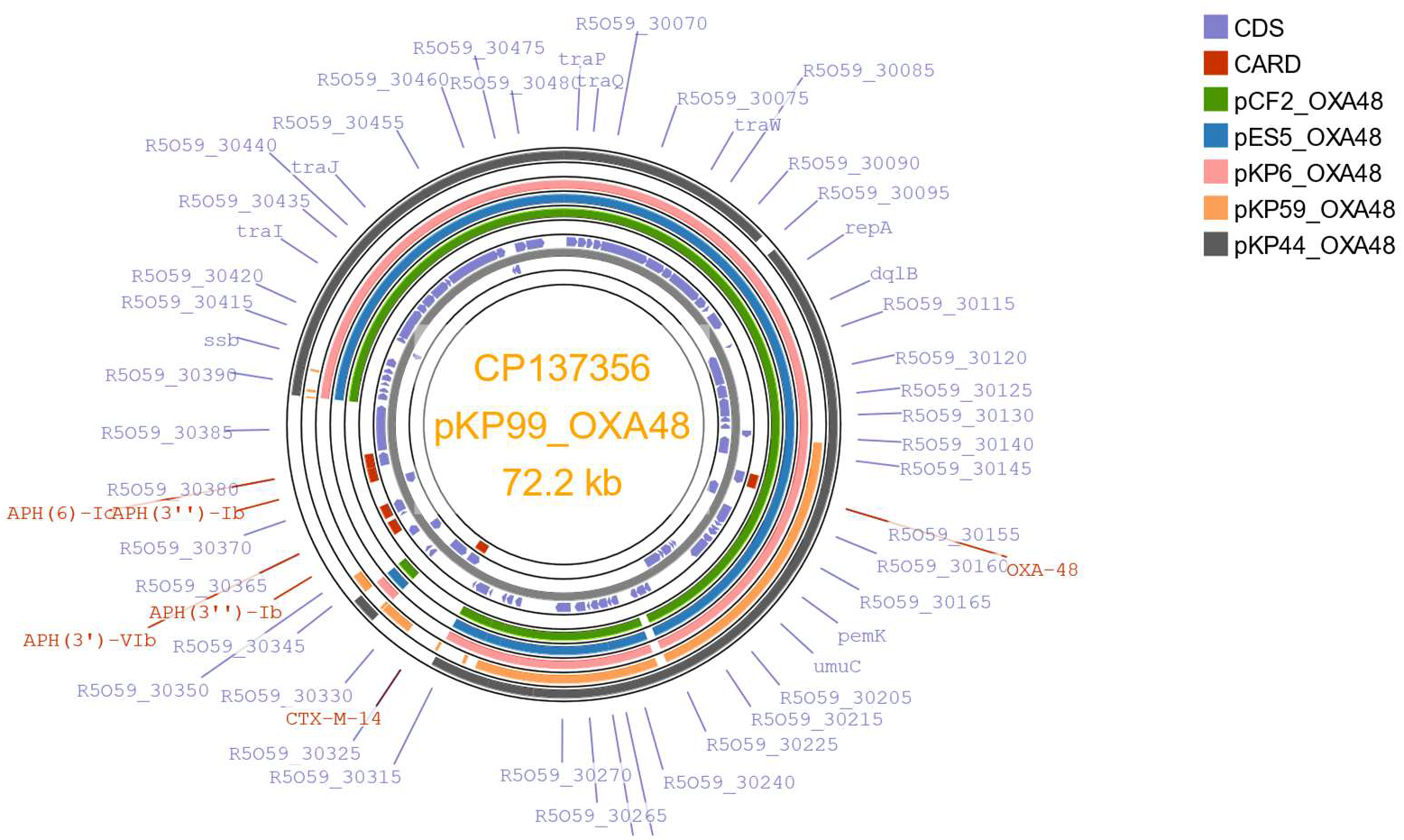
BLAST comparison of plasmids carrying *bla*_OXA-48_. Coloured regions show areas of homology. Most isolates carried IncL plasmids of approximately 64 kb (e.g. pCF2_OXA48, pES5_OXA48 and pKP59_OXA48), while pKP99_OXA48 had an extra segment carrying additional resistance genes. KP44_NW4_44.22_ST307_OXA48 had an IncM1 plasmid of 61.5 kb (pKP44_OXA48). The IncR plasmid pKP59_OXA48 (54.8 kb) had only partial homology with the IncL/M plasmids. Isolates compared were from four different hospitals, with pCF2_OXA48, pES5_OXA48 and pKP59_OXA48 being from isolates from the same hospital group. Plasmids were compared using Proksee [8]. Resistance genes were identified by the Comprehensive Antibiotic Resistance Database (CARD) on Proksee. CDS, coding sequences.

In agreement with our results, Boyd et al [12] also commented in a recent review that *bla*_OXA-48_ is usually found in self-conjugative 60- to 70-kb plasmids with an IncL scaffold, but that there is some variation among them, with some plasmids also carrying other resistance genes. Notably, the *bla*_OXA-48_ gene is not associated with class 1 integrons in these plasmids, and is typically carried in transposon Tn*1999* with an upstream, and often a downstream, IS*1999* insertion sequence [13].

### OXA-181 plasmids

The set included nine isolates carrying *bla*_OXA-181_ from six different hospitals. Most carried ColKP3 replicons (e.g. <u>CP137409</u> in KP102_WM3_17.23_ST147_OXA181, <u>CP141549</u> in KP75_L14_02.23_ST20_OXA181 and <u>CP141544</u> in KP89_L16_06.23_ST307_OXA181) and either no, or only *qnrS1* resistance gene in addition. The 51,479 bp ColKP3/IncX3 plasmids pKP75_OXA181 and pKP89_OXA181 matched (100 % identity, 100 % coverage) one another and many others on GenBank (e.g. <u>CP034284</u>, <u>MK412917</u>, <u>MK412920</u>, <u>CP114995</u> and <u>CP110780</u>) from isolates of *K. pneumoniae, E. coli*, the *Enterobacter cloacae* complex and *Citrobacter freundii* from various countries (e.g. USA, United Arab Emirates, Italy) (Fig. 3). However, *bla*_OXA-181_ was associated with IncFII replicon types in some isolates (e.g. KP4_L4_52.21_ST147_OXA181). Li *et al*. [14] who studied 191 *bla*_OXA-48-like_ plasmids in *K. pneumoniae* also noted that *bla* _OXA-181_ was often carried by 50-kb ColKP3-IncX3 conjugative plasmids, which were found in multiple countries, and in small ColKP3-type mobilizable plasmids, mirroring our results.

**Fig. 3.**
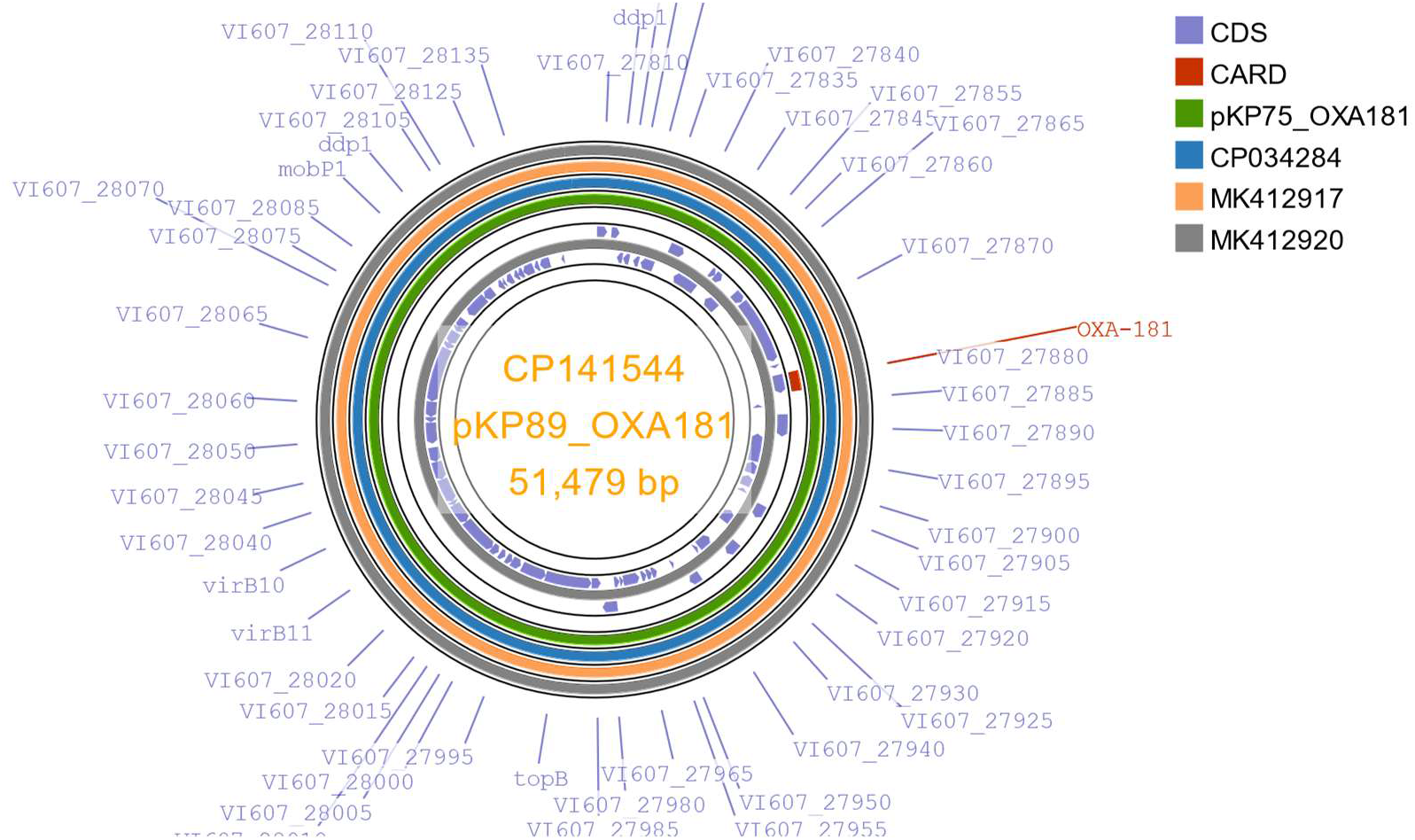
BLAST comparison of 51.5 kb ColKP3/IncX3 plasmids from isolates of *K. pneumoniae* from two different hospitals and belonging to different sequence types from this study with examples from GenBank from isolates of *K. pneumoniae* (<u>CP034284</u> and <u>MK412920</u>) and *E. coli* (<u>MK412917</u>) from USA and United Arab Emirates. All the plasmids matched one another (100 % identity, 100 % coverage). CDS, coding sequences; CARD, Comprehensive Antibiotic Resistance Database

In common with our results finding identical ColKP3/IncX3 plasmids carrying *bla*_OXA-181_ from isolates of different types and genera and from various countries, others have also observed that *bla*_OXA-181_ are predominantly associated with IncX3 and ColKP3 plasmids and that these are near identical among isolates from Jordan, Egypt, Turkey, South Korea, Thailand, South Africa, Kuwait, China and Angola [15,16].

### OXA-232 plasmids

The set included nine isolates carrying *bla*_OXA-232_ from seven different hospitals. As with the *bla*_OXA-181_ plasmids, those carrying *bla*_OXA-232_ were often associated with ColKP3 replicons (e.g. <u>CP140296</u>, <u>CP141573</u>, <u>CP137419</u>, <u>CP141552</u>). They carried no other resistance genes. The original assemblies indicated that three isolates carried a ColKP3 plasmid of 6.14 kb (KP28_L6_38.22_ST147_NDM5_OXA232, KP67_L13_49.22_ST437_NDM5_OXA232 (<u>CP141552</u>) and KP104_L5_18.23_ST147_NDM5_OXA232), 5 carried 12.3 kb plasmids (KP21_L5_33.22_ST16_NDM5_OXA232,, KP33_NE2_41.22_ST2096_OXA232 (<u>CP140296</u>), KP35_L8_42.22_ST2096_OXA232 (<u>CP141573</u>), KP54_WM4_48.22_ST147_OXA232 and KP101_SE2_14.23_ST147_NDM5_OXA232 (<u>CP137419</u>)) and one a 18.4 kb plasmid (KP24_L5_34.22_ST16_NDM5_OXA232), but closer examination showed that the 12.3 kb plasmids consisted of 2 copies of the 6.14 kb plasmid and the 18.4 kb plasmid 3 copies, perhaps because of an assembly issue with flye with high copy number reads. Analysis of the reads themselves showed that they mostly exist as 6.14 kb single copy plasmids although they may also be present as double and triple copy plasmids. There were many matches/close matches to these plasmids on GenBank (e.g. <u>CP01562</u>, <u>JX423831</u>, <u>OY856406</u>). The original description of OXA-232 described the gene in 6.1 kb plasmids [17] (e.g. <u>JX423831</u>), but there are also examples on GenBank of the 12.3 kb double version (e.g. in FDAARGOS_440 <u>CP023924</u>). Li *et al*. [14] also commented that the *bl*a_OXA-232_ gene was mainly found in small (6.1-kb) ColKP3-type plasmids, largely found in India in their study.

### KPC-2 plasmids

The majority of the isolates carrying *bla*_KPC-2_ in this study were from hospital WM3 and have been described earlier; they carried mainly IncN or IncHI2/ IncHI2A *bla*_KPC-2_ plasmids (Fig. 1). The set also included five isolates from a further two hospital groups. Those from hospital WM4 all belonged to *K. pneumoniae* ST307 and all carried a 52.3 kb IncN plasmid containing *bla*_KPC-2_ and *bla*_TEM-1B_ (e.g. <u>CP133741</u>). They lacked the segment carrying *aph(6)-Id, aph(3”)-Ib, qnrB2, sul1* and *dfrA19* found in the IncN plasmids from hospital WM3. That from hospital YH1 carried a 99.3 kb IncR/IncN plasmid with *bla*_KPC-2_, *bla*_TEM-1A_, *bla*_OXA-9_ and *dfrA14* (<u>CP133867</u>).

### KPC-3 plasmids

Four isolates in the set from two hospital groups carried plasmids with *bla*_KPC-3_. All belonged to *K. pneumoniae* ST307 and their *bla*_KPC-3_ plasmids were 90 to 97 kb in size with those from hospital L9 carrying *bla*_TEM-1A_ and *bla*_OXA-9_ in addition, while those from hospital WM1 carried no additional resistance genes (e.g. KP19_WM1_28.22_ST307_KPC3). All were of IncFIB(pQil)/IncFII(K) replicon types. The closest matches on GenBank were slightly bigger plasmids (109 to 118 kb) that shared 99 % identity and 99 % coverage with pKP19_KPC3 (e.g. <u>OQ821101</u>).

### IMP-1 plasmids

The set included 13 isolates carrying *bla*_IMP-1_ from six different hospitals, with six isolates belonging to *K. pneumoniae* novel sequence type ST6601 from 2 hospitals (L1 and L6) between which patient transfers are known to have occurred. These isolates carried IncHI2/IncHI2A plasmids of approximately 270 kb carrying *bla*_IMP-1_, *aac(6’)-Ib3, bla*_SHV-12_, *sul1* and *qnrA1*. Isolates KP20_YH2_29.22_ST54_IMP1, EB1_SW1_27.22_ST104_IMP1, EB2_SW1_42.22_ST145_IMP1 and EB3_SW1_43.22_ST134_IMP1 all carried IncHI1A(NDM-CIT)/IncHI1B(pNDM-CIT) plasmids of 334 to 361 kb in size carrying *bla*_IMP-1_, *aac(3)-IIa, aac(6’)-Ib3, bla*_DHA-1_, *mph(A), qnrB4, sul1, dfrA17* (e.g. <u>CP141538</u> in EB3_SW1_43.22_ST134_IMP1), despite being isolated from two different regions of the UK and all belonging to different types. Interestingly, the plasmids from the EB2 and KP20 isolates lacked the copper and silver heavy metal resistance genes found in the plasmid from the EB3 isolate (Fig. 5). KO2_WM3_48.22_STNV2_IMP1 carried a 170.5 kb IncC plasmid containing *bla*_IMP-1_, *aac(6’)-Ib-cr, aac(6’)-Ib3, qnrA1, sul1* and dfrA1, while KP40_L1_43.22_ST20_IMP1 had a 57.0 kb IncN3 plasmid containing *bla*_IMP-1_, *aac(6’)-Ib3, qnrA1* and *sul1* (<u>CP141568</u> in KP40_L1_43.22_ST20_IMP1), which was highly similar to those we have reported previously from *Enterobacter* isolates from London hospitals from 2019 (<u>CP043856</u> and <u>CP043516</u>) [18] (Fig. 6). *Acinetobacter baumannii* isolate AB1_L10_11.23_ST203_IMP1 carried *bla*_IMP-1_, *aadA5, aac(6’)-IIa, msr(E), mph(E), floR, sul1* and *sul2* in a 326.9 kb plasmid with no plasmid hits on PlasmidFinder.

**Fig. 4:**
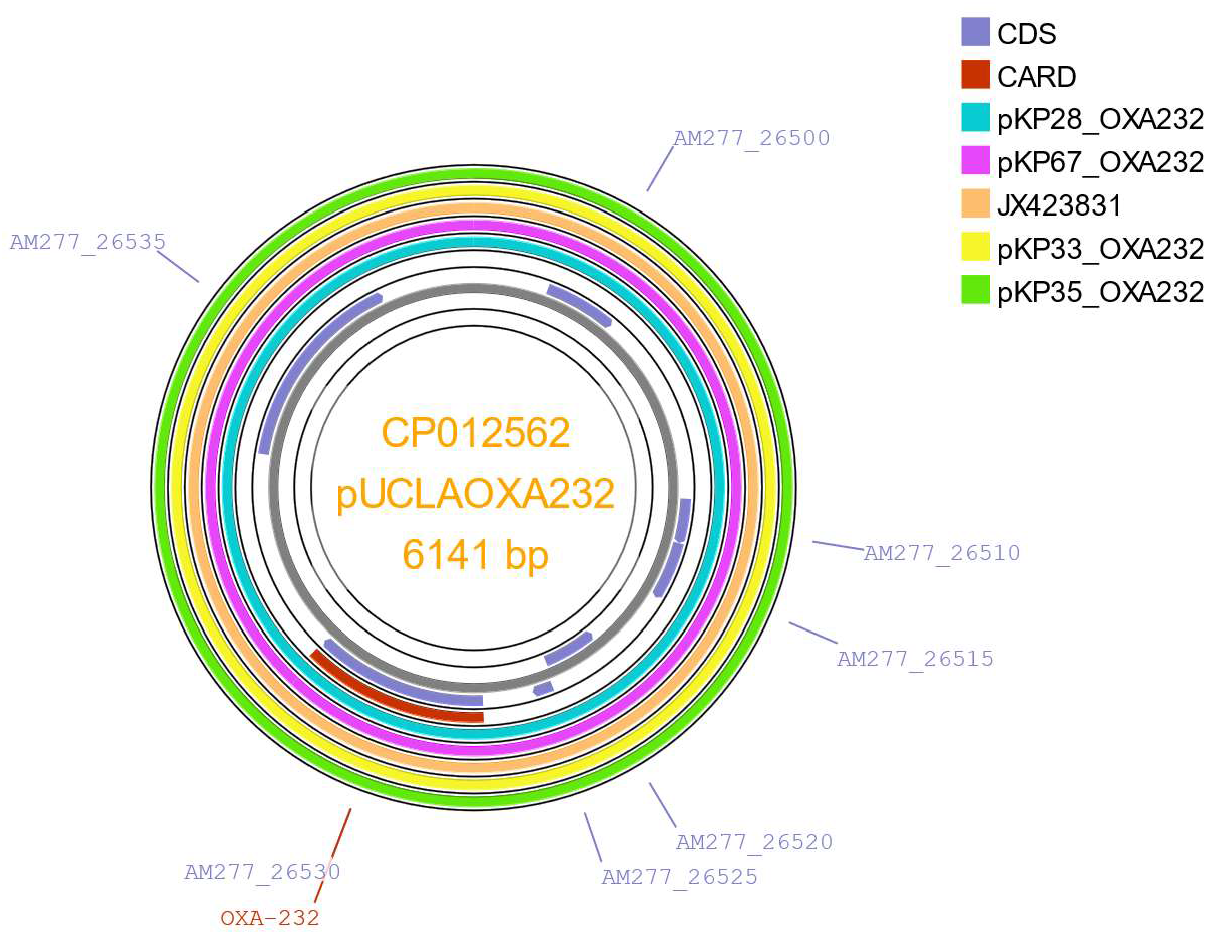
BLAST comparison of ColKP3 plasmids carrying *bla*_OXA-232_ in isolates of *K. pneumoniae* from this study with <u>CP012562</u> (pUCLAOXA232-1) from an isolate of *K. pneumoniae* from California, USA and <u>JX423831</u> from an isolate of *E. coli* from France (transferred from India). Plasmids compared are from KP28_L6_38.22_ST147_NDM5_OXA232, KP33_NE2_41.22_ST2096_OXA232, KP35_L8_42.22_ST2096_OXA232 and KP67_L13_49.22_ST437_NDM5_OXA232, which are from four different hospitals. Plasmids were compared using Proksee [8]. Resistance genes were identified by the Comprehensive Antibiotic Resistance Database (CARD) on Proksee. CDS, coding sequences.

**Fig. 5.**
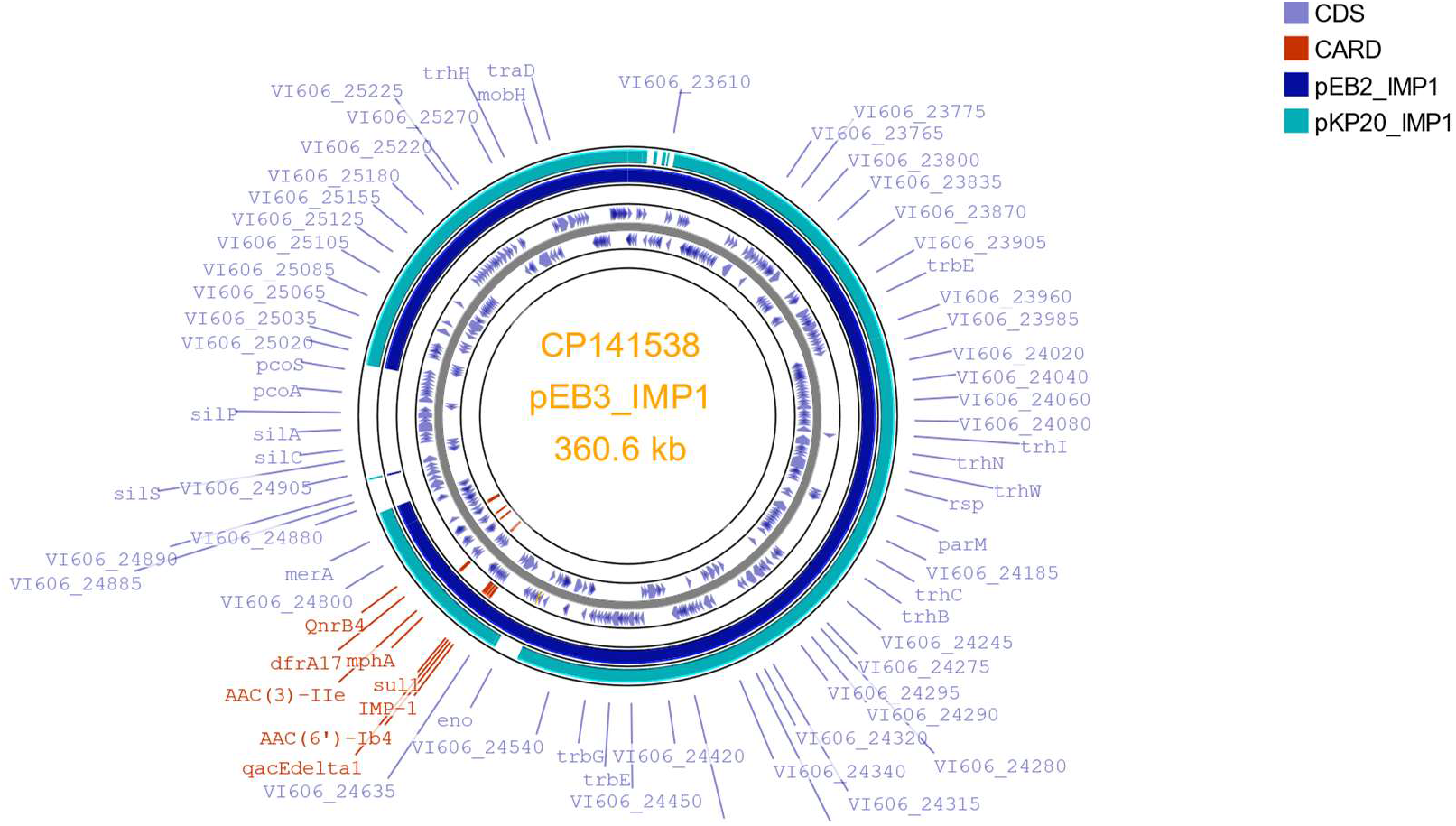
BLAST comparison of the IncHI1A(NDM-CIT)/IncHI1B(pNDM-CIT) plasmids carrying *bla*_IMP-1_ from *E. cloacae* complex and *K. pneumoniae* isolates from two different hospitals (pEB2_IMP1 and pEB3_IMP1 from hospital SW1, pKP20_IMP1 from hospital YH2). The *Enterobacter* isolates belonged to distinct sequence types (STs 145 and 134). CDS, coding sequences; CARD, Comprehensive Antibiotic Resistance Database.

**Fig. 6.**
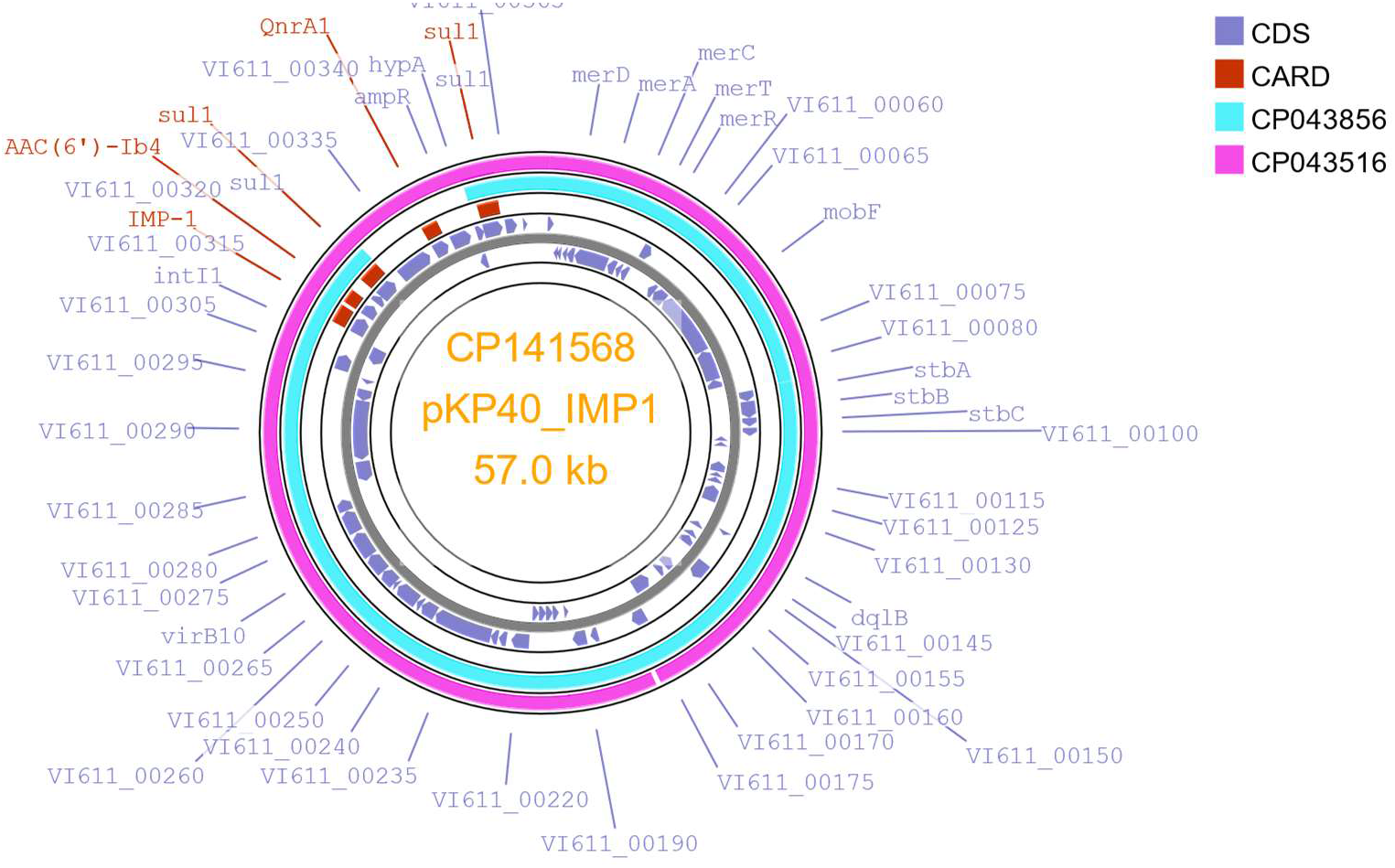
BLAST comparison of the IncN3 plasmid from *K. pneumoniae* KP40_L1_43.22_ST20_IMP1 with IncN3 plasmids from *Enterobacter* isolates from London hospitals from 2019 (<u>CP043856</u> and <u>CP043516</u>). CDS, coding sequences; CARD, Comprehensive Antibiotic Resistance Database,

### IMP-4 plasmids

The set included just two isolates carrying *bla*_IMP-4_ which were both from the same hospital and belonged to the same type (KP12_NE1_13.22_ST54_IMP4 and KP14_NE1_17.22_ST54_IMP4). Both had IncHI2/IncHI2A plasmids of approximately 326 kb containing *bla*_IMP-4_, *aph(3”)-Ib, aac(6’)-Ib3, aac(6’)-Ib-cr, aph(6)-Id, mcr-9, bla*_SHV-12_, *bla*_TEM-1B_, *bla*_OXA-1_, *mph(A), qnrB2, sul1, dfrA19, ARR-3, formA*, truncated *qacE* and *catB3*. Although not well-represented in this set, we are aware that large IncHI2/IncHI2A plasmids carrying both *bla*_IMP-4_ and *mcr-9* are found in an increasing proportion of IMP positive isolates in the UK [19], with most isolates referred to the reference laboratory during the latter part of 2023 carrying them. Such plasmids are widely distributed and have been described in other organisms and countries/continents (e.g. <u>KX810825</u> in *Salmonella enterica* MU1, <u>CP022533</u> in *Enterobacter hormaechei* <u>MS7884A</u> both from Australia) [20] (Fig. 7).

**Fig. 7.**
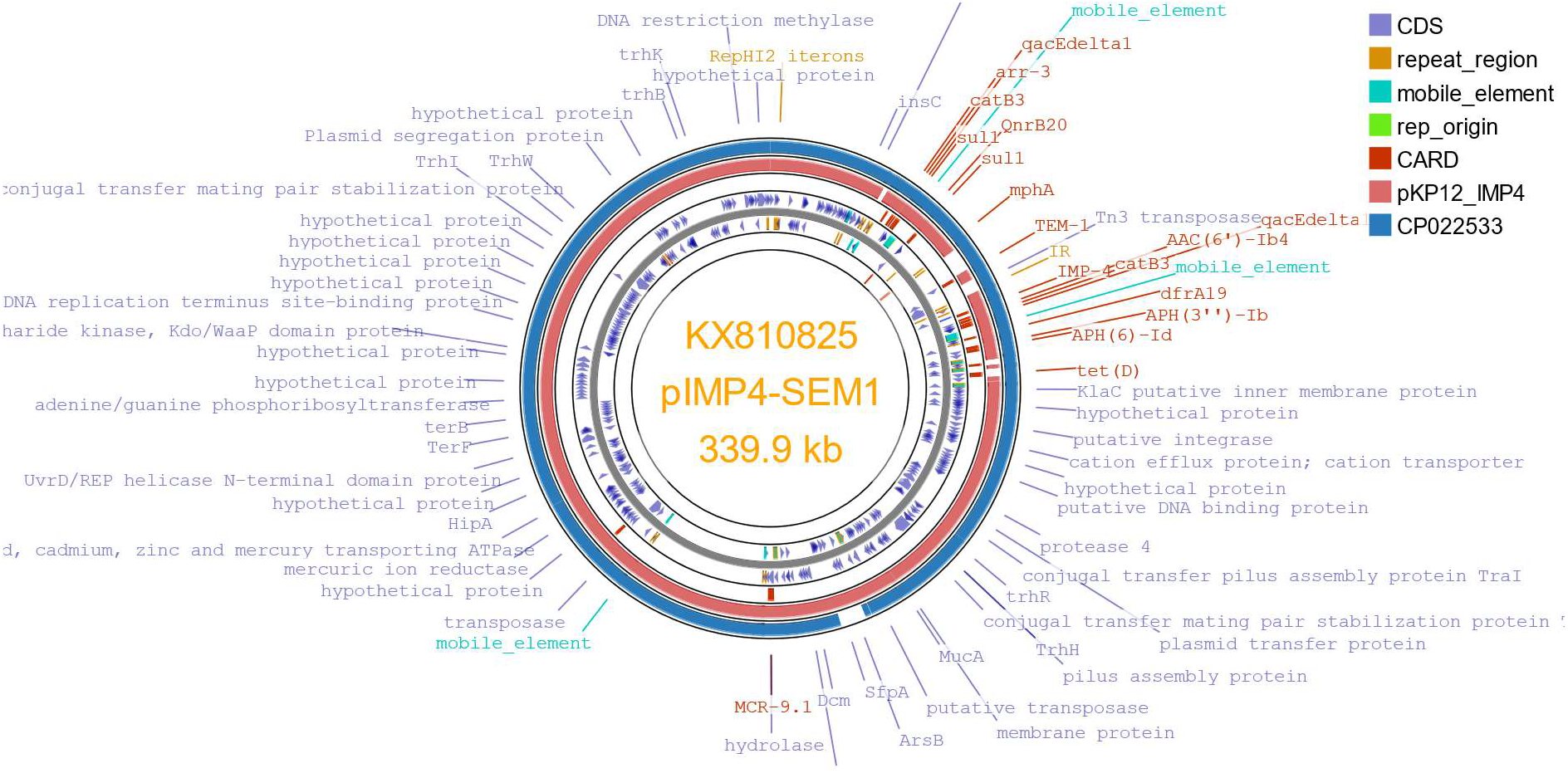
BLAST comparison of the IncHI2/IncHI2A *bla*_IMP-4_ plasmid in *K. pneumoniae* KP12_NE1_13.22_ST54_IMP4 with those from *Salmonella enterica* MU1 (<u>KX810825</u>) and *Enterobacter hormaechei* MS7884A (<u>CP022533</u>) from Australia. CDS, coding sequences; CARD, Comprehensive Antibiotic Resistance Database.

### NDM-1 plasmids

Six of the 12 isolates with *bla*_NDM-1_ sequenced were representatives of *K. pneumoniae* ST20 from hospital L5 described earlier which had IncHI2/IncHI2A plasmids of approximately 340 kb containing *bla*_NDM-1_, *aadA5, aac(6’)-Ib-cr, aac(6’)-IIc, bla*_DHA-1_, *bla*_OXA-1_, *ere(A), mph(A), catB3, qnrB4, sul1* and *dfrA17* (e.g. <u>CP141854</u>). KP118_NW3_20.23_ST101_NDM1 similarly carried an IncHI2/IncHI2A *bla*_NDM-1_ plasmid, which was slightly larger (365.7 kb) and contained *aph(6)-Id, aph(3”)-Ib, bla*_SHV-12_, *bla*_TEM-1B_ and *catA2* resistance genes in addition to *bla*_NDM-1_. A further isolate (KP36_NW3_42.22_ST834_NDM1) had a 131.2 kb IncFIB(pQil) plasmid with *bla*_NDM-1_, *aac(3)-IIa, aac(6’)-Ib-cr, bla*_CTX-M-15_, *bla*_TEM-1B_, *bla*_OXA-1_, truncated *catB3* and *qnrB1*. The remaining isolates were representatives of *K. pneumoniae* ST147, three of which (from two different hospitals) had 54 kb IncFIB(pQil) plasmids with *bla*_NDM-1_, *bla*_CTX-M-15_, *bla*_OXA-1_, *ARR-3, sul1, qnrS1, aac(6’)-Ib-cr, aph(3’)-VI, catB3* and truncated *qacE* (e.g. <u>CP137381</u>). The remaining representative of ST147 from a further hospital had a 352.3 kb IncFIB(pNDM-Mar)/IncHI1B(pNDM-MAR) hybrid virulence resistance plasmid carrying *bla*_NDM-1_, *aph(3’)-Ia, aph(3’)-VI, armA, msr(E), mph(A), mph(E), qnrS1, sul1, sul2* and *dfrA5* and virulence plasmid elements aerobactin (*iutA, iucABCD*), *rmpA*/*A2, terABCDEWXYZ*, hemin, *luxR, pagO, shiF*, lysozyme inhibitor (<u>CP137400</u>).

### NDM-5 plasmids

The rise in frequency of this NDM variant, particularly in *E. coli*, but also in *K. pneumoniae*, has been marked and has been noted by many authors [9, 10, 11]. Our set included two representatives of *E. coli* ST167 with multi-replicon IncF plasmids carrying *bla*_NDM-5_ (e.g.<u>CP133853</u>) along with *aac(6’)-Ib-cr, aadA2, sul1, dfrA12, sitABCD, tet(A), bla*_CTX-M-15_, *bla*_OXA-1_, *mph(A)*, truncated *qacE* and *catB3* (ES3_WM2_48.22_ST167_NDM5 and ES4_WM2_48.22_ST167_NDM5). Other isolates carrying *bla*_NDM-5_ included in this study belonged largely to *K. pneumoniae* ST147 (or ST6796, a single locus variant of ST147) (n=15), which was a particular focus and are described more fully elsewhere [21]. All but five of these carried an IncFIB(pNDM-Mar)/IncHI1B(pNDM-MAR) hybrid virulence resistance plasmid that combined *bla*_NDM-5_ and other resistance genes with genes associated with virulence plasmids (aerobactin (*iutA, iucABCD*), *rmpA*/*A2, terABCDEWXYZ*, hemin, *shiF*, lysozyme inhibitor) (e.g. <u>CP137386</u> in KP39_L11_43.22_ST147_NDM5, <u>CP137375</u> in KP58_WM2_48.22_ST147_NDM5_OXA48, CP137372 in KP81_L15_03.23_ST147_NDM5, <u>CP137435</u> in KP82_L15_04.23_ST147_NDM5, <u>CP137360</u> in KP97_L17_12.23_ST147_NDM5, <u>CP137352</u> in KP99_L14_14.23_ST147_NDM5_OXA48 and <u>CP137392</u> in KP124_L5_21.23_ST147_NDM5), despite being from seven different hospital groups. The remaining five representatives of ST147 did not have a virulence plasmid and carried *bla*_NDM-5_ in 96 kb IncFII plasmids also with *aadA2, rmtB, bla*_TEM-1B_, truncated *qacE, erm(B), dfrA12, sul1* and *mph(A)* (KP2_L3_37.21_ST147_NDM5 and KP30_YH2_39.22_ST147_NDM5_OXA181), or in a 156.5 kb IncFII/IncR plasmid also carrying *aadA2, aadA1, armA, rmtB, bla*_CTX-M-15_, *bla*_TEM-1B_, *msr(E), ere(A), mph(A), mph(E), erm(B), cmlA1, qnrB1, ARR-2, sul1* and *dfrA12* (KP104_L5_18.23_ST147_NDM5_OXA232) or chromosomally (KP29_L6_38.22_ST147_NDM5 and KP28_L6_38.22_ST147_NDM5_OXA232), the latter result being obtained on two separate occasions for both isolates.

Five further isolates of *bla*_NDM-5_ positive *K. pneumoniae* belonging to four further STs were included; the two isolates belonging to ST16 (both from hospital L5) had a 46.1 kb IncX3 plasmid carrying no other resistance gene, while that of ST3096, also from hospital L5, carried *bla*_NDM-5_ in a 143 kb IncC plasmid that also contained *aac(6’)-Ib3, bla*_CMY-23_ and *sul1*. That belonging to ST1558 (KP77_WM1_02.23_ST1558_NDM5) had *bla*_NDM-5_ in a 348 kb IncFIB(pNDM-Mar)/IncHI1B(pNDM-MAR) hybrid virulence resistance plasmid also carrying *aph(3’)-Ia, aadA1, aph(3’)-VI, aac(6’)-Ib, bla*_CTX-M-15_, *bla*_TEM-1C/1B_, *bla*_OXA-9_, *mph(A)*, truncated *catA1, sul1* and *dfrA5*, aerobactin genes, *rmpA*/*A2*, tellurite resistance genes, *shiF*, lysozyme inhibitor and *cobW* (<u>CP141546</u>) that was highly similar to those seen in representatives of ST147, with most similarity to <u>CP137375</u> from KP58_WM2_48.22_ST147_NDM5_OXA48. Similar plasmids have also been seen in *K. pneumoniae* ST383 (e.g. <u>CP034201</u> and <u>CP091814</u>) and it is worrying that they have been seen in multiple types and among geographically unrelated isolates. Finally, that belonging to ST437, from a further hospital (L13) had a 106.8 kb IncFIB(pQil)/IncFII plasmid carrying *bla*_NDM-5_ together with *bla*_CTX-M-15_, *erm(B), sul1* and *dfrA30*.

### NDM-14 plasmids

Just three isolates in the set, which were clonally and epidemiologically related, carried this unusual NDM allele (KP86-KP88 _L3_06.23_ST147_NDM14). All belonged to *K. pneumoniae* ST147 and carried a 54.1 kb IncFIB(pQil) plasmid containing *bla*_NDM-14_, *aph(3’)-VI, aac(6’)-Ib-cr, bla*_CTX-M-15_, *bla*_OXA-1_, *catB3, qnrS1, ARR-3* and *sul1* (e.g. <u>CP137365</u>, <u>CP137425</u>). All also carried a separate hybrid virulence/resistance plasmid (e.g. <u>CP137363</u>, <u>CP137424</u>). There are a number of highly similar plasmids in isolates of *K. pneumoniae* on GenBank of the same length (99-100 % coverage, 99.99-100 % identity) e.g. <u>CP021947</u>, <u>CP014757</u>, <u>OQ785270</u>, <u>CP014757</u> and <u>AP018834</u> from isolates from the USA, South Korea and Myanmar, although most carried *bla*_NDM-1_ rather than *bla*_NDM-14_.

### Virulence/resistance plasmids in *K. pneumoniae*

In addition to the hybrid virulence resistance plasmids described in representatives of ST147 (and single locus variant (SLV) ST6796) [21], we have described hybrid virulence resistance plasmids in other, less common types in this study; ST1558 (<u>CP141546</u>) (KP77_WM1_02.23_ST1558_NDM5), ST2096 (<u>CP140295</u>) (KP33_NE2_41.22_ST2096_OXA232) (an SLV of ST14) and ST395 (<u>CP133748</u>) (KP17_NW3_25.22_ST395_OXA48 and KP59_WM2_48.22_ST395_OXA48). In the case of the latter three isolates, these plasmids did not include a carbapenemase gene, which was carried in another plasmid in both cases. In common with other non-NDM hybrid resistance virulence plasmids found in representatives of ST147 they were large (approximately 310 kb) IncFIB(pNDM-Mar)/ IncHI1B(pNDM-MAR) plasmids carrying *armA, msrE, mphE, sul2* and *catA1* (KP33) or *armA, aadA2, bla*_CTX-M-15_, *bla*_TEM1A_, *msrE, dfrA12, sul1, tetD, mphE* (KP17) in addition to aerobactin genes, *rmpA*/*rmpA2*, tellurite resistance genes, *shiF*, lysozyme inhibitor and *cobW*. That of KP77_WM1_02.23_ST1558_NDM5 carried *bla*_NDM-5_ and was highly similar to the NDM-5 virulence resistance plasmids found in representatives of ST147 (e.g. <u>CP137375</u>).

### Carbapenemase genes among representatives of *K. pneumoniae* ST307

*K. pneumoniae* ST307 is a ‘high-risk’ clone for resistance that is commonly received by the reference laboratory from hospitals and has been used as an exemplar for comparison of plasmids found within the same type within and between hospitals. We typed 144 non-duplicate isolates of this type from 2021-2023 of which at least 32 were received as carbapenemase producers.

The set included 22 representatives of ST307 from nine hospital laboratories and four regions carrying a variety of carbapenemase genes; *bla*_KPC-2_ (n=10), *bla*_KPC-3_ (n=4), *bla*_OXA-181_ (n=2) and *bla*_OXA-48_ (n=6). Largely, the plasmids found were consistent within a hospital set and distinct from those from other hospitals carrying the same carbapenemase gene; for example, the 3 KPC-positive isolates from hospital WM4 (KP68_WM4_50.22_ST307_KPC2, KP72_WM4_01.23_ST307_KPC2 and KP73_WM4_01.23_ST307_KPC2) all had a 52.3 kb IncN plasmid carrying *bla*_KPC-2_ and *bla*_TEM-1B_, while that from hospital YH1 carried *bla*_KPC-2_ in a 99.3 kb IncR/IncN plasmid with *bla*_TEM-1A_, *bla*_OXA-9_ and *dfrA14*. Another set of isolates from hospital NW4 from five patients all carried *bla*_OXA-48_ in 61.5 kb IncM1 plasmids with no other resistance genes, and all carried an additional IncFII(K)/IncFIB(K) plasmid with *aac(3)-IIa, aph(6)-Id, aph(3”)-Ib, aac(6’)-Ib-cr, bla*_CTX-M-15_, *bla*_TEM-1B_, *bla*_OXA-1_, truncated *catB3, qnrB1, sul2, tetA* and *dfrA14*, supporting that they are linked. Both isolates from hospital WM1 with *bla*_KPC-3_ carried 92.9 kb IncFIB(pQil)/IncFII(K) plasmids.

These results are in contrast with those of representatives of ST147, many of which shared highly similar hybrid virulence plasmids despite coming from different hospital groups.

### Further Comments and Conclusions

It is clear that comparison of plasmids to investigate if ‘plasmid outbreaks’ are occurring is far from straightforward. Even where similar plasmids were found among isolates that were related in time and space there were often minor differences between them with segments missing/additional segments (e.g. Fig. 1) leading to potential difficulties in interpretation. In addition, many plasmids are widely distributed, and finding similar plasmids among isolates carrying them does not necessarily indicate plasmid spread between them. Examples are the widespread IncL pOXA48a and IncHI2/IncHI2A *bla*IMP-4 plasmids that have been found in many countries [19, 20, 22]. The widely found ColKP3/IncX3 OXA-181 plasmids and ColKP3 *bla*_OXA-232_ plasmids were remarkably conserved (to the base) and seen in different organisms and many different geographical locations. It is clearly important to know the context and wider distribution of a relevant plasmid before considering whether a plasmid outbreak may be occurring. All three hospital sets we examined revealed a complex picture. In all of them, the situation was mixed with some evidence for plasmid transfer, but also clear examples where the plasmids were different.

IncHI2/HI2A plasmids are important vehicles for carbapenemase genes; we observed plasmids of this type carrying *bla*_KPC-2_, *bla*_IMP-1_, *bla*_IMP-4_ or *bla*_NDM-1_. Where we observed the colistin resistance gene *mcr-9* it was always in IncHI2/IncHI2A plasmids e.g. <u>CP141854</u>, although not all carried this gene. It is likely that the gene is carried silently and is not expressed in these isolates [23].

This study concentrated on the plasmids carrying carbapenemase genes, but most of these isolates carried multiple plasmids with the others often also carrying resistance genes. For example, not only did KP72_WM4_01.23_ST307_KPC2 carry a KPC plasmid (<u>CP141579</u>), but it also had a further plasmid containing *aac(3)-IIa, aph(6)-Id, aph(3”)-Ib, aac(6’)-Ib-cr, bla*_CTX-M-15_, *bla*_TEM-1B_, *bla*_OXA-1_, truncated *catB3, qnrB1, sul2, tet(A)* and *dfrA14* (<u>CP141580</u>). This study also only looked at these plasmids at a relatively superficial level and a deeper analysis could achieve greater insight.

While most plasmids carrying *bla*_OXA-48_ did so in a typical IncL plasmid of approximately 60-70 kb, there was some variation among these plasmids and some very different plasmids were found (e.g. the IncR plasmid pKP59_OXA48), showing that there can be value in analysing plasmids carrying *bla*_OXA-48_ where a plasmid outbreak is suspected.

Nanopore sequencing was essential in providing complete or near complete assemblies of these isolates, facilitating investigation of their plasmids. Wide use of nanopore sequencing is providing information not previously available leading to a greatly increased understanding of the global epidemiology of plasmids carrying carbapenemase genes.

---

Table S1. Carbapenemase producers included in the study and their resistance genes. Isolates were labelled with a unique organism number (beginning KP for *K. pneumoniae*, EB for *E. cloacae* complex, ES for *E. coli*, CF for *C. freundii*, KO for *K. oxytoca* complex and AB for *Acinetobacter baumannii*), followed by a hospital code, the week and year of receipt by the laboratory, their sequence type (ST) and the carbapenemase genes that they carried. Hospitals were labelled according to region in the UK and number within that region (L, London; SE, South East England; SW, South West England; WM, West Midlands; EM, East Midlands; NE, North East London; NW, North West England; YH, Yorkshire and Humber). Contigs are labelled as circular (c) or linear (l). An asterisk indicates a truncated gene or (in the allelic profile column) a novel allele. Where there was no exact match to a resistance gene, the closest alternative(s) are given. NV refers to novel sequence types. Where more than one run was carried out on an isolate, run identifications highlighted in turquoise indicate that the same extract was used; yellow highlighting indicates that a different extract was used.

